# Surface phase separation in a confined space

**DOI:** 10.64898/2025.12.03.692075

**Authors:** Daoning Wu, Jie Lin

## Abstract

Biomolecular condensates frequently form near surfaces, including lipid membranes. However, the effect of a surface on phase separation within a cell, a confined system, remains elusive. In this work, we study surface phase separation under the constraint of a finite number of molecules in a confined space and identify three wetting states: thin, intermediate, and thick. Notably, the intermediate state, which is unstable in a fixed-chemical-potential system, can be stabilized in confinement below a critical width. Intriguingly, in wider systems, the intermediate solution becomes unstable due to a negative relationship between the chemical potential and the particle number, resulting in partial-wetting or prewetting droplets. Importantly, starting from a homogeneous surface, droplets can form only when the average volume fraction reaches the threshold of the intermediate state. Our theory elucidates the kinetic pathways of droplet formation in a confined system, providing crucial insights into how biomolecular condensates near surfaces respond to cellular events, such as the synthesis of biomolecules.

## Introduction

Numerous experiments have shown that liquid-liquid phase separation (LLPS) drives the formation of biomolecular condensates that are essential for various cellular functions [1–10]. To date, theoretical studies of LLPS have primarily focused on bulk systems, where surface effects are not considered. However, cells are full of surfaces: plasma membranes, organelle membranes, nuclear membranes, etc [11–13]. Many experiments have reported critical interaction between membranes and membraneless organelles, e.g., regulation of membraneless organelles by endoplasmic-reticulum [14, 15], formation of autophagosomes at the surface of condensates [16], scission of endosome membranes by biomolecular condensates [17], and even the formation of transcriptional condensates on the surface of DNA [18].

Simulations and theories have suggested that these surfaces constrain molecular mobility and amplify local concentrations through surface phase separation [19–22]. In a binary solution where a dense and a dilute phase co-exist, a surface that has a high affinity with the solute molecules exhibits a wetting transition at the wetting-transition temperature *T*_*w*_ [23–26]. Typically, the wetting transition is a first-order transition between a partial wetting state (a thin layer covering the surface, potentially with wetting droplets) and a complete wetting state (a thick layer covering the entire surface).

The classical theory of surface phase separation developed by Cahn considered a semi-infinite system with a fixed chemical potential [23]. In this framework, the grand potential is minimized at thermal equilibrium, and the number of molecules is not conserved. However, cell compartments such as the cytoplasm and the nucleus are confined spaces. We expect that thermodynamic equilibrium can be established below the time scale when molecule synthesis and degradation are significant, supported by the observations that P granule assembly and disassembly are governed by phase separation based on local thermal equilibrium [6]. Indeed, for bulk phase separation, most theoretical studies and even textbooks have focused on systems with a fixed number of molecules [4, 27–30]. A biologically relevant theory of surface phase separation should also study a confined system with a fixed number of molecules. Are the classical theories still valid, and are there any unexpected but biologically/physically critical effects in a confined system? To our knowledge, these questions are still elusive.

Here, we address these questions by studying a binary solution in a confined space that interacts with an attractive surface. First, we investigate the stability and configuration of homogeneous wetting, where the volume fraction of solute molecules depends only on the distance *z* from the surface. We reveal three homogeneous wetting solutions: thin, intermediate, and thick. Remarkably, in the intermediate state, an increase in the number of molecules leads to a decrease in chemical potential, indicating that it is thermodynamically unstable [31]; however, it can remain stable in a confined system with a fixed number of molecules. Surprisingly, we unveil a new critical temperature *T*_*sc*_, distinct from the prewetting temperature *T*_*pw*_ and the wetting temperature *T*_*w*_ of the classical theory of wetting transition [23, 25]. Above *T*_*sc*_, the system exhibits a continuous change of the surface wetting profile as one changes the total number of solute molecules. In contrast, below *T*_*sc*_, the system exhibits a discontinuous transition between two states with different thicknesses.

We next allow the volume fraction profile to be inhomogeneous in the *x* direction parallel to the surface. Using linear stability analysis, we demonstrate that there exists a threshold system width *d*_*c*_, above which the intermediate state is unstable against small perturbations. In particular, 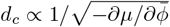, where *µ* is the chemical potential and 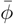 is the average volume fraction. Below the wetting temperature, the intermediate state evolves into a partial-wetting state, characterized by droplets with a finite contact angle. Above the wetting temperature, the intermediate state evolves into a prewetting state characterized by pancake-like droplets of finite thickness. Remarkably, the instability of the intermediate state itself is independent of whether the temperature *T* is below or above the wetting temperature *T*_*w*_. We confirm all of our predictions by simulating a two-dimensional continuum system.

Our theory sheds light on how biomolecular condensates near surfaces transition between different wetting configurations in response to cellular triggers, such as the synthesis or degradation of molecules, and to changes in physiological parameters, including temperature, pH, and molecular crowding. In recent experiments by Morin et al. [18], the authors observed a transition from a thin absorbed layer of transcription factors to a partial-wetting state in which multiple droplets wet the DNA as the total number of transcription factors increased. We remark that this partial-wetting state is likely triggered by the transition between the thin and intermediate solutions, as a consequence of increasing 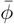 in our phase diagram below *T*_*w*_ (Figure 3a). To confirm this, we also simulate a quasi-static increase or decrease of 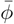 in a two-dimensional system. Indeed, when 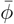 increases to the intermediate region, droplets start to form on the surface, and the system state is history dependent because both the homogeneous states and partial-wetting/prewetting states can be locally stable.

### The model

We study a binary solution in a confined space where the system length in the *z* direction is *L*. To simplify the problem, we let the volume fraction of the solute molecule *ϕ* be homogeneous in the *xy*-plane and only depend on *z*. Later, we will extend our model to allow the volume fraction to vary along the *x* direction. The system’s Helmholtz free energy per unit area is

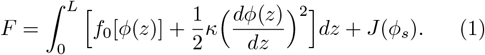

Here, *f*_0_[*ϕ*(*z*)] is the free energy density, 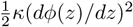 is the energy cost due to the gradient of *ϕ*(*z*) [32], and *J*(*ϕ*_*s*_) is the interaction energy between the surface at *z* = 0 and the solute molecule where *ϕ*_*s*_ ≡ *ϕ*(*z* = 0). For simplicity, we use the free energy of a regular solution [28]:

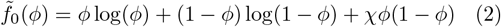

Here, 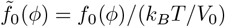 where *f*_0_ is the Helmholtz free energy density, *k*_*B*_ is the Boltzmann constant, *T* is the temperature, and *V*_0_ is the molecule volume. The bulk interaction strength *χ* = 2*T*_*c*_*/T*, where *T*_*c*_ is the bulk critical temperature. When *T < T*_*c*_, the system exhibits phase separation in bulk with a dense phase and a dilute phase. We set the interaction energy between the surface and the solute molecules as *J*(*ϕ*_*s*_) = − *aϕ*_*s*_ where *a >* 0. In the following, we use the non-dimensionalized version of the model unless otherwise specified: the energy density unit is *ϵ*_0_ = *k*_*B*_*T/V*_0_ and the length unit is 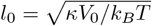.

For a system with a fixed number of molecules, 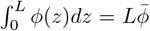 is constant, and the Helmholtz free energy is minimized in equilibrium. In contrast, for a system with a fixed chemical potential, the grand potential

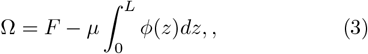

is minimized in equilibrium. To study the volume fraction profile for a system of a fixed number of molecules at thermal equilibrium, a trick is to compute the equilibrium profile and the resulting 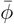 given a fixed chemical potential *µ* and then find the appropriate *µ* that gives the target 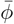. As we show later, this method enables us to find all possible equilibrium states that minimize the Helmholtz free energy.

Minimization of the grand potential leads to the equilibrium condition as 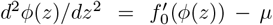 for 0 *< z < L* with the boundary condition *ϕ*^*′*^(*z*) = −*a* at *z* = 0 and *ϕ*^*′*^(*z*) = 0 at *z* = *L*. The boundary condition at *z* = *L* results from the absence of interaction between the surface at *z* = *L* and the solute molecules. The equilibrium condition leads to the conservation equation *ϕ*^*′*^(*z*)^2^*/*2 = *W* (*ϕ*(*z*)) where we introduce the W function *W* = *f*_0_(*ϕ*) − *µϕ* − *f*_0_(*ϕ*_*L*_) + *µϕ*_*L*_. The constant terms in *W* allow the boundary condition *ϕ*^*′*^(*z* = *L*) = 0 to be satisfied. In this work, we mostly study an attractive surface, and the volume fraction monotonically decreases from the surface at *z* = 0, which means that

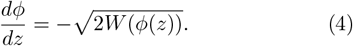

We treat the chemical potential in *W* (*ϕ*) as a changing parameter, and the intersection of the line −*J*^*′*^(*ϕ*) and 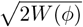 determines the equilibrium surface volume fraction *ϕ*_*s*_ (Figure 1a-c). Given a fixed chemical potential, there are three possible solutions of *ϕ*_*s*_ at most. The solution below *ϕ*_*L*_ is not considered because the volume fraction near an attractive surface must be higher than *ϕ*_*L*_. We denote the three solutions from low to high value of *ϕ*_*s*_ as thin, intermediate, and thick (Figure 1d-f). As we show later, the intermediate solution can be stable for a confined system; in contrast, it must be unstable for a system with a fixed chemical potential [25]. We remark that the chemical potential can be lower (Figure 1a-b) or higher (Figure 1c) than the equilibrium value *µ*_*c*_ at which bulk phase separation occurs (for the regular-solution model, *µ*_*c*_ = 0). The thick solution for *µ > µ*_*c*_ is unphysical and excluded because *ϕ*^*′*^(*z*) is imaginary when *W <* 0 (Figure 1c).

**FIG. 1.**
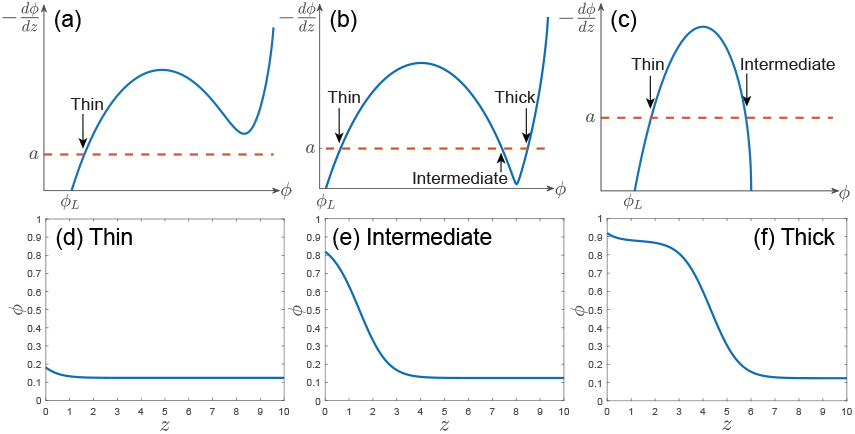
(a-c) The intersections between −*J*^*′*^(*ϕ*) (the red dashed line) and 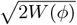 (the blue solid line) determine the volume fraction at the surface, *ϕ*_*s*_. Here, we show three different chemical potentials with one (a), three (b), and two (c) solutions. (d-f) The equilibrium volume-fraction profiles for the thin (d), intermediate (e), and thick (f) solution. In this work, we set *a* = 0.1 and *L* = 10 unless otherwise mentioned.

Given the chemical potential, we compute the average volume fraction 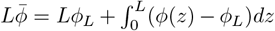, which we rewrite as

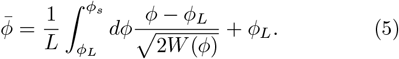

The chemical potential is related to *ϕ*_*L*_ by 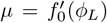 since near *z* = *L, ϕ*(*z*) is virtually a constant, which we confirm numerically later. We prove that there is always a one-to-one correspondence between *µ* and *ϕ*_*L*_ (Supplemental Material).

### Homogeneous equilibrium profiles

We compute the phase diagram of wetting transition in a confined space as a function of the average volume fraction of solute molecules and temperature (Figure 3a). We remark that for a system with a fixed molecule number, the emergence of three homogeneous solutions occurs at the prewetting critical temperature *T*_*pw*_ [23, 25]. Above *T*_*pw*_ (the orange dashed line in Figure 3a), the intermediate solution does not exist. The thin and thick solutions are indistinguishable from each other. We also highlight the wetting temperature *T*_*w*_ (the green dashed line in Figure 3a) at which point the thick and thin solutions have the same grand potential if the system’s chemical potential is fixed at the bulk-phase-separation value *µ*_*c*_ [23]: below *T*_*w*_ at *µ*_*c*_, droplets, if formed, must have finite contact angles; above *T*_*w*_ at *µ*_*c*_, droplets with finite contact angles cannot form.

Surprisingly, we unveil a previously unknown critical temperature at *T*_*sc*_ (the purple dashed line in Figure 3a) that is unique for systems with a fixed molecule number below which a bistable region emerges (Figure 3a). For *T > T*_*sc*_, as one continuously changes 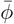, a unique solution exists (Figure 2a, b). The transition between the thin and intermedi ate solutions occurs at 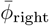, where −*J*^*′*^(*ϕ*) is tangent to 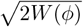 at its local maximum (Figure 2a inset). The transition point between the intermediate and thick solutions occurs at 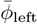, where −*J*^*′*^(*ϕ*) is tangent to 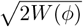 at its local minimum (Figure 2a inset). As 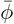 quasi-statically decreases from high to low, the volume fraction profile transitions through thick, intermediate, and thin solutions sequentially, with a smooth transition in between. We verify our predictions numerically by directly minimizing the Helmholtz free energy with a conserved number of molecules with a homogeneous volume-fraction profile *ϕ*(*z*). We confirm that the equilibrium profiles are at the minimum point of the Helmholtz free energy (Figure 2b inset). We note that the chemical potential at 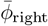 is higher than *µ*_*c*_ in Figure 2. Nevertheless, if the temperature is close enough to the bulk critical value, the chemical potential at 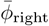 can still be lower than *µ*_*c*_ (Figure S2).

**FIG. 2.**
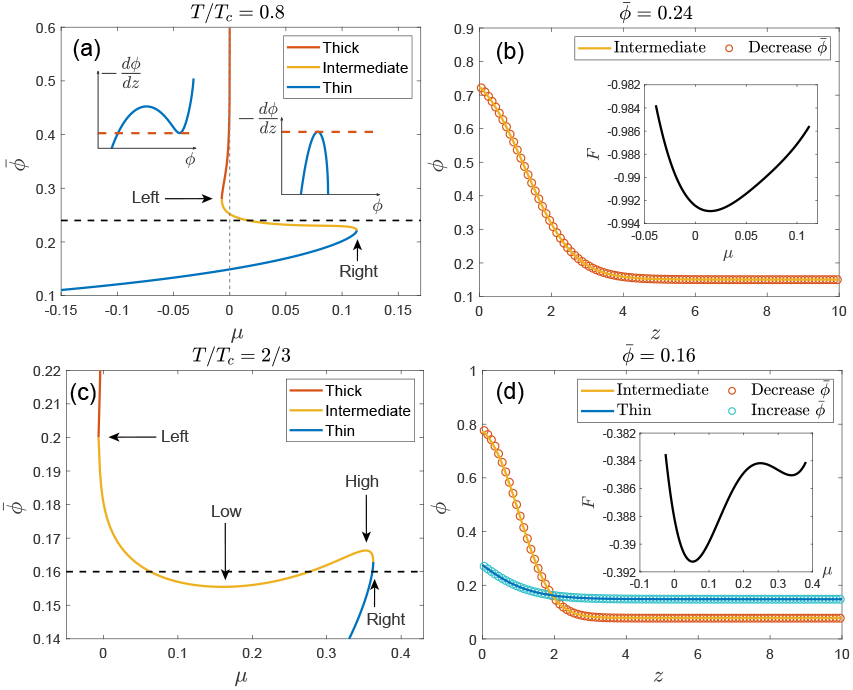
(a) 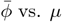 for *T > T*_*sc*_. The insets show *J*^*′*^(*ϕ*) (the red dashed line) and 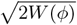 (the blue solid line) at 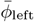 and 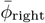. (b) The theoretical (solid line) and numerical (circles) volume-fraction profiles at 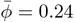 for an intermediate solution. The inset shows the Helmholtz free energy *F* as a function of *µ*. (c) 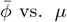 for *T > T*_*sc*_. (d) The theoretical and numerical volume-fraction profiles at 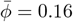, where the volume-fraction profile is history-dependent. If 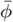 decreases from 0.17 to 0.16, the profile takes the intermediate solution; in contrast, if 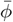 increases from 0.15 to 0.16, the profile takes the thin solution. The inset shows the Helmholtz free energy *F* as a function of *µ* with two minima.

Intriguingly, as 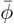 decreases from high to low, the chemical potential exhibits a non-monotonic behavior, which increases during the intermediate state (Figure 2a). Remarkably, the intermediate solution is stable even if 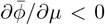. For bulk systems, a necessary condition for a stable state is 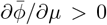; otherwise, the system is unstable against random fluxes of molecules [31]. In our case, an attractive surface in a confined space breaks down the stability condition of bulk systems. Later, we show that the intermediate solution indeed becomes unstable when the system width in the *x* direction exceeds a threshold.

In contrast, for *T < T*_*sc*_, a first-order transition occurs: as 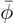 decreases from high to low and reaches 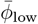, the intermediate solution disappears. The system abruptly jumps to a thin solution (Figure 2c). In contrast, as 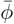 increases from low to high and reaches 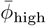, the system abruptly jumps from one intermediate solution to another with lower *µ* (Figure 2c). We verify the existence of hysteresis numerically by implementing a quasi-static decrease or increase of 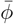. As predicted, the equilibrium profile depends on the history of parameter changes (Figure 2d), and the Helmholtz free energy also exhibits two minima in the bistable region (Figure 2d inset).

We remark that as *L* approaches infinity, the first term on the right of Eq. (5) can be neglected for the intermediate solutions. Therefore, we must have 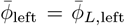 and 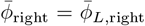, which means that 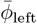 approaches the bulk saturation concentration in the *L*→ ∞ limit at low temperature (Figure S3, S4).

### Inhomogeneous profiles

We next allow the volume-fraction profile to be inhomogeneous in the *x* direction. Imagine introducing a small perturbation *δh*(*x, t*) to a flat interface whose initial height is *h*_0_. The surface tension tends to restore the interface to flatness. Meanwhile, we can imagine cutting the system into many small compartments, each with a small width. A local increase in the interface height in one compartment increases its local 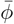, which leads to a decrease in its local chemical potential if 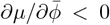, inducing a molecule flux into the compartment and further amplifying the height increment. Combining the two competing factors, we propose the following dynamics of *δh*(*x, t*) in the linear order:

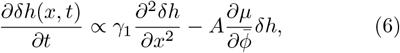

Here, *γ*_1_ is the surface tension constant, and *A* is a constant depending on the diffusion constant of the solute molecules. By introducing a periodic perturbation to the interface, *δh*(*x, t*) = *h*_1_ cos(*qx*), which satisfies 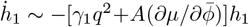, we identify the critical wavelength 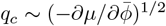 below which, the system is unstable against perturbation. Therefore, when the system width *d* is above *d*_*c*_, which satisfies

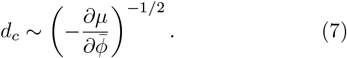

Interfacial instability occurs for the intermediate solution.

We numerically simulate a two-dimensional system in which the molecular flux is proportional to the gradient of the chemical potential (see simulation details in the Supplemental Material). Starting from an intermediate homogeneous state, we add a small perturbation to the profile and observe the resulting interface shape. We confirm the existence of a minimum system width to generate the interfacial instability and its relation to the derivative between *µ* and 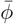 (Figure 3b). As we predict, the thick and thin states are always stable (Figure S5).

**FIG. 3.**
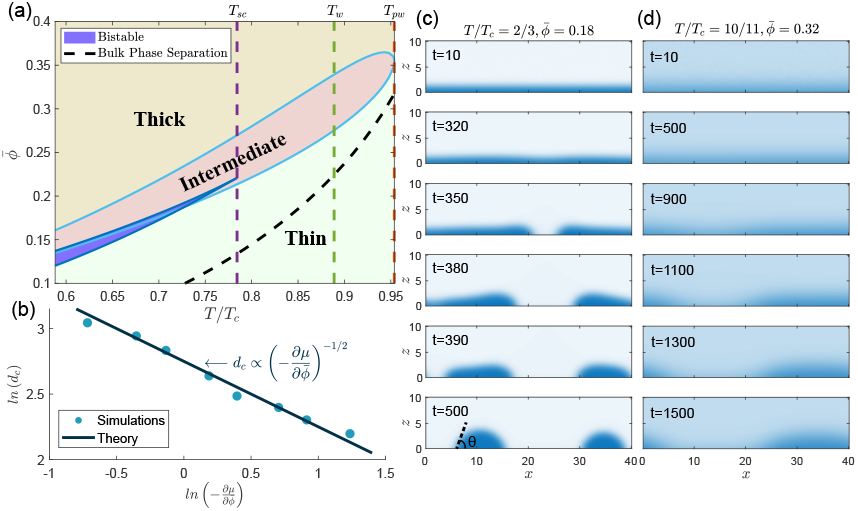
(a) The phase diagram of surface phase separation in a confined system with a fixed number of molecules. The bistable region can be divided into an upper region with two coexisting intermediate solutions and a lower region with an intermediate solution and a thin solution that coexist. The black-dashed line shows the saturation concentration for bulk phase separation. (b) Log-log plot of the minimum width to generate instability 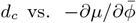. The simulation results for *T/T*_*c*_ = 2*/*3 fall approximately on a line of slope -1/2, verifying our prediction. (c) Simulations of the two-dimensional system below the wetting temperature *T*_*w*_ at different times. Periodic boundary conditions are applied in the *x* direction with *d* = 40. At *t* = 0, a small perturbation is added to a homogeneous intermediate profile. The measured contact angle *θ* ≈ 70^°^, close to *θ*_*theory*_ = 72.7^°^ predicted by Young’s formula. (d) The same as (c) but for *T > T*_*w*_ .

What might be the final states after the initial instability of the intermediate solution? Intriguingly, the unstable intermediate state evolves to a partial-wetting state where the chemical potential approximately equals the bulk-phase-separation value *µ*_*c*_ for *T < T*_*w*_ (Figure 3c). In this case, the measured contact angle of the finite droplets are close to the value predicted by Young’s formula, which can be well approximated as *γ* cos(*θ*) = *aϕ*_*b*_ − *aϕ*_*a*_ (Figure 3c), where *γ* is the surface tension between the dense and dilute phase, and *ϕ*_*a*_ and *ϕ*_*b*_ are the volume fractions of the dense phase and the dilute phase, respectively (Supplemental Material) [21]. Meanwhile, for *T > T*_*w*_, the droplets in the final state have blurred boundaries and pancake-like shapes (Figure 3d), corresponding to the prewetting state where the thin and thick states coexist. Indeed, the volume-fraction profiles of the droplet region and the non-droplet region, as a function of *z*, are close to the predicted profiles of the thick solution and the thin solution, respectively (Figure S6), given that the chemical potential is fixed at the simulated value. We note that the intermediate state and the thin state can be bistable for *T < T*_*sc*_ for a homogeneous system. In this case, we find that the initial intermediate state also evolves into a partial-wetting state, rather than the thin state (Figure S7).

To investigate how the synthesis or degradation of molecules affects wetting configuration, we simulate a quasi-static increase of 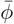 from a low value (Figure 4a, c, and Movie S1, S2). When 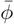 reaches the threshold of the intermediate state, which is much higher than the bulk saturation volume fraction (Figure 3a, even in the *L* limit, see the 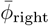 curve in Figure S4), partial-wetting or prewetting droplets form quickly on the surface as long as the instability condition is satisfied. As 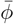 keeps increasing, droplets grow in size until they fully occupy the system, which then transitions to the homogeneous thick state (not shown in Figure 4a).

**FIG. 4.**
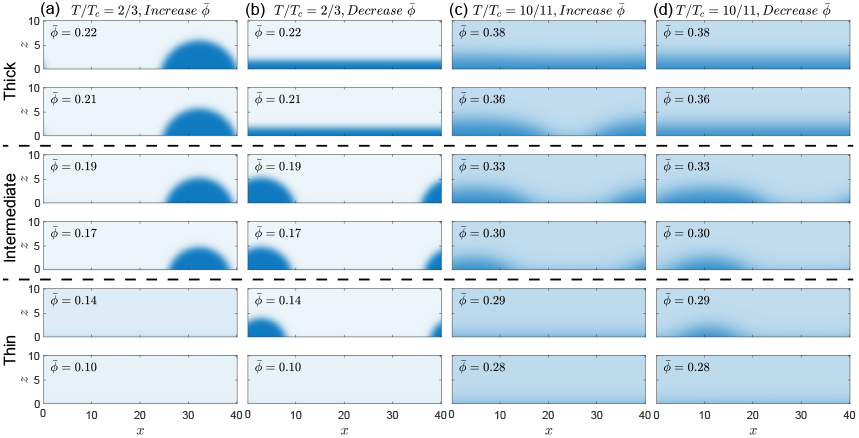
Simulated volume fraction distributions when 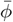 increases (a,c) or decreases (b,d) quasi-statically, below (a,b) or above (c,d) the wetting temperature. We highlight the entrance and departure of the intermediate state by the dashed lines.

When decreasing 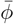 from a high value, the initial homogeneous layer can be maintained until the system reaches the intermediate region (Figure 4b, d and Movie S3, S4). Again, the partial wetting/prewetting state is locally stable and can be maintained even in the thin region. We notice that when 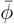 exceeds the critical value for spinodal decomposition, droplets can form in the bulk. However, this state is unstable since the molecules will be absorbed by the surface (Movie S5).

## Discussion

In this work, we study the wetting transition in a confined space with a fixed number of molecules. Compared with the traditional Cahn model, where the chemical potential is fixed, our model is more relevant to surface separation phenomena in biological systems. To our surprise, the intermediate solution, which is deemed unstable in the fixed-chemical-potential scenario, can be stable in our case as long as the system width is below a threshold. Furthermore, we identify a new critical temperature *T*_*sc*_, below which the transition of volume-fraction profiles is discontinuous.

When the system width is above the threshold value *d*_*c*_, the intermediate solution is unstable against infinitesimal perturbations and tends to evolve to a partial-wetting (*T < T*_*w*_) or prewetting state (*T > T*_*w*_), depending on the temperature. Importantly, as one quasistatically increases the average volume fraction 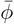 from a low value, the homogeneous state becomes unstable only when 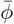 reaches the threshold of the intermediate state, which can be much higher than the bulk saturation volume fraction. Interestingly, the partial wetting/prewetting states are locally stable, and the system’s actual state, whether homogeneous in the *x* direction or not, depends on its history.

The interfacial instability identified in this work is different from other instability mechanisms. For example, Rayleigh-Taylor instability is generated due to competition between gravity and surface tension; spinodal dewetting is triggered by attractive long-range forces [24]. To our knowledge, interfacial instability resulting from a negative correlation between chemical potential and particle number has not been previously reported.

We thank Shengyao Luo, Lingyu Meng, and Xueping Zhao for helpful discussion related to this work. The research was funded by the National Key Research and Development Program of China (2024YFA0919600), National Natural Science Foundation of China (Grant No. 12474190) and Peking-Tsinghua Center for Life Sciences grants.

