## Supplemental Material for "Surface phase separation in a confined space"

Daoning Wu[1] and Jie Lin[2,3]
[1] *Yuanpei College, Peking University, Beijing, China*
[2] *Center for Quantitative Biology, Academy for Advanced Interdisciplinary Studies, Peking University, Beijing, China*
[3] *Peking-Tsinghua Center for Life Sciences, Academy for Advanced Interdisciplinary Studies, Peking University, Beijing, China*
(Dated: December 3, 2025)

#### Proof of the one-to-one correspondence between $\mu$ and $\phi_L$

In the main text, we introduce the W function $W(\phi) = f_0(\phi) - \mu\phi - f_0(\phi_L) + \mu\phi_L$. Because $W(\phi_L) = 0$ and $W'(\phi_L) = 0$, we find that

$$W(\phi_L + \delta\phi) = \frac{1}{2}W''(\phi_L)(\delta\phi)^2 + O((\delta\phi)^3) = \frac{1}{2}f_0''(\phi_L)(\delta\phi)^2 + O((\delta\phi)^3), \tag{S1}$$

where $0 < \delta\phi \ll 1$. Whenever there is a solution for the equilibrium volume-fraction profile, $W(\phi_L + \delta\phi)$ should be positive; otherwise, $\phi'(z) = -\sqrt{2W(\phi(z))}$ is imaginary. Thus, we conclude that $f_0''(\phi_L) > 0$; in other words, $\mu = f_0'(\phi_L)$ increases monotonically with $\phi_L$.

#### Details of numerical simulations

In simulations of homogeneous profiles, the system is one-dimensional in the $z$ direction. We divide the system into $N$ cells along the $z$ direction, each with a uniform width of $\Delta z = L/N$, where $L$ is the system length in the $z$ direction. The volume fraction in the $i$-th cell is $\phi_i$ ($i = 1, 2, ..., N$). Considering that $\mu = f_0'(\phi) - d^2\phi/dz^2$, the chemical potential of the cells are

$$\mu_i = f_0'(\phi_i) - \frac{\phi_{i+1} + \phi_{i-1} - 2\phi_i}{(\Delta z)^2}, \qquad i = 1, 2, ..., N. \tag{S2}$$

$\phi_0$ and $\phi_{N+1}$ are the volume fractions in the auxiliary cells (the cells with dashed borders in Figure S1). To enforce the boundary condition $\phi'(z) = -a$ at $z = 0$ and $\phi'(z) = 0$ at $z = L$, we let $\phi_0 = \phi_1 + a\Delta z$ and $\phi_{N+1} = \phi_N$ after each time step.

Let $J_i$ be the flux of solute molecule from the $i$-th cell to the $i+1$-th cell. As the system is bounded at $z = 0$ and $z = L$, we have $J_0 = J_N = 0$. The remaining fluxes can be calculated by

$$J_i = \frac{\mu_i - \mu_{i+1}}{\Delta z}, \qquad i = 1, 2, ..., N-1. \tag{S3}$$

For each time step $\Delta t$, we update the volume fractions with

$$\Delta\phi_i = (J_{i-1} - J_i)\Delta t, \qquad i = 1, 2, ..., N. \tag{S4}$$

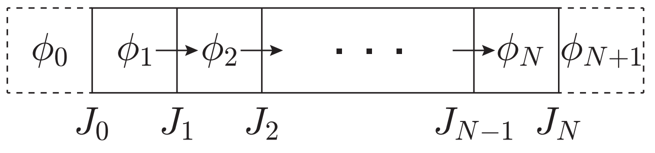


FIG. S1. A schematic of the numerical simulations in one dimension.

In simulations of two-dimensional inhomogeneous profiles, the system is divided into cells with width $\Delta z = L/N$ in both the $x$ and $z$ directions. The approach to the boundary condition in the $z$ direction is the same as the homogeneous case. For the $x$ direction, a periodic boundary condition is applied with auxiliary cells.

In this work, a small perturbation is added by superimposing a random number matrix with a sum of zero onto the volume fraction distribution, where the random numbers are uniformly distributed between -0.01 and 0.01.

### Estimation of contact angle with Young's formula

In the main text, we approximate Young's formula as $\gamma\cos(\theta) = a\phi_b - a\phi_a$, where $\theta$ is the predicted contact angle and $\gamma$ is the surface tension between the dense and dilute phases. We use the volume fractions of the dense phase and the dilute phase to approximately compute the solid-dilute-phase surface tension $a\phi_a$ and the solid-dense-phase surface tension $a\phi_b$, respectively. To compute the surface tension between the dilute and dense phase, we consider an infinite system in which the volume fraction only depends on the coordinate $z$ and satisfies $\phi(z=\infty) = \phi_a$ and $\phi(z=-\infty) = \phi_b$. The system's chemical potential is fixed at the bulk-phase-separation value, and its grand potential per unit area is

$$\Omega = \int_{-\infty}^{\infty} \Big[f_0(\phi) + \frac{1}{2}\Big(\frac{d\phi}{dz}\Big)^2 - \mu_c\phi - f_0(\phi_a) + \mu_c\phi_a\Big]dz. \tag{S5}$$

Minimizing the grand potential leads to the equilibrium condition $d^2\phi(z)/dz^2 = f_0'(\phi(z)) - \mu_c$. The equilibrium condition leads to the conservation equation

$$\frac{1}{2}\Big(\frac{d\phi(z)}{dz}\Big)^2 = W(\phi) \tag{S6}$$

where the W function here is $W(\phi) = f_0(\phi) - \mu_c\phi - f_0(\phi_a) + \mu_c\phi_a$. Since the volume fraction must decrease from $z=-\infty$ to $z=\infty$, we obtain

$$\frac{d\phi}{dz} = -\sqrt{2W(\phi)}. \tag{S7}$$

Finally, we obtain the expression of the surface tension constant, which is simply the grand potential per unit area as

$$\gamma = \int_{-\infty}^{\infty}\Big[\frac{1}{2}\Big(\frac{d\phi(z)}{dz}\Big)^2 + W(\phi)\Big]dz = \int_{-\infty}^{\infty}\Big(\frac{d\phi(z)}{dz}\Big)^2 dz = \int_{-\infty}^{\infty}\frac{d\phi(z)}{dz}d\phi = \int_{\phi_a}^{\phi_b}\sqrt{2W(\phi)}d\phi. \tag{S8}$$

For $T/T_c = 2/3$, $\phi_a = 0.0707$, $\phi_b = 0.9293$, from which we can calculate that $\gamma = 0.2883$. Thus, the contact angle predicted by the Young's formula is $\theta_{theory} = 72.7°$.

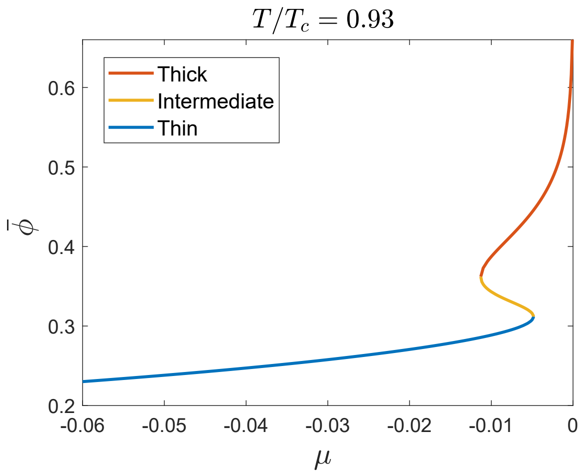


FIG. S2. $\bar{\phi}$ vs. $\mu$ for the case where $\sqrt{2W(1/2)} < a$ when $\mu = \mu_c$. In this case, the chemical potential is always lower than $\mu_c$ as $\bar{\phi}$ changes.

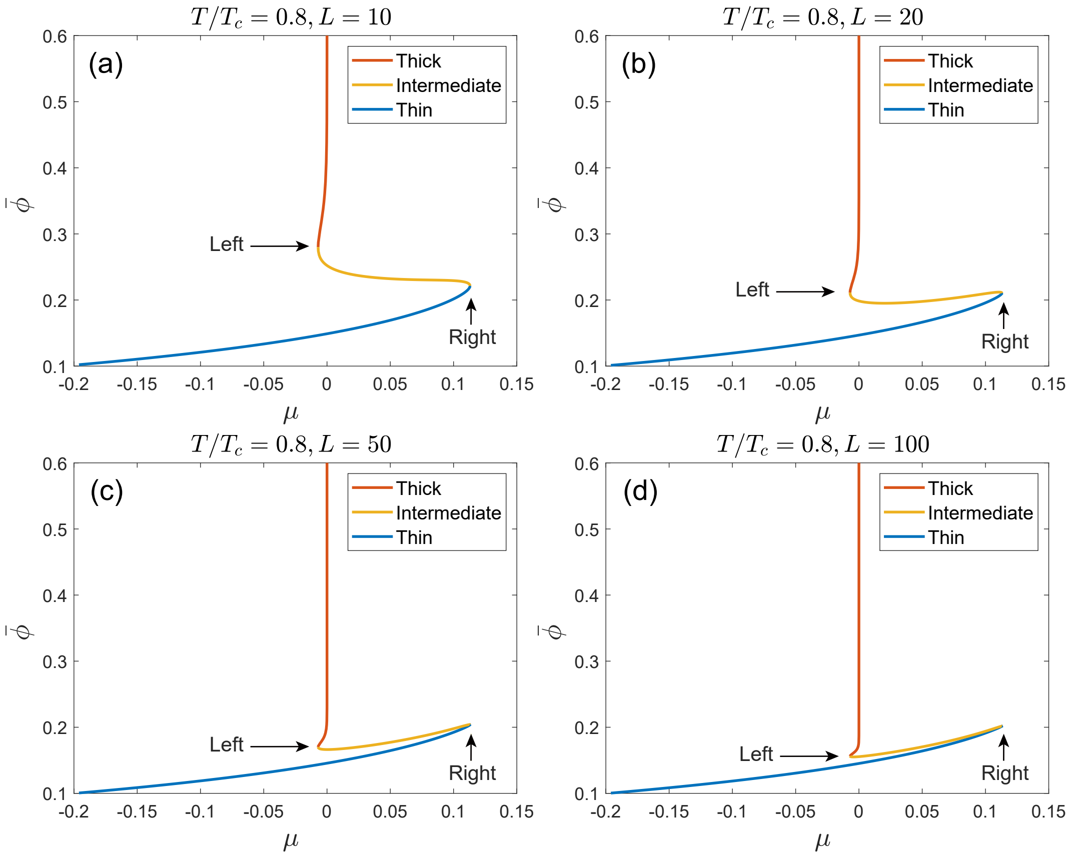


FIG. S3. The change of $\bar{\phi}$ vs. $\mu$ as $L$ increases from 10 to 100. The other parameters are the same as those in Figure 2a of the main text. One should note that $\bar{\phi}_{\text{right}}$ is larger than $\bar{\phi}_{\text{left}}$ when $L$ is large.

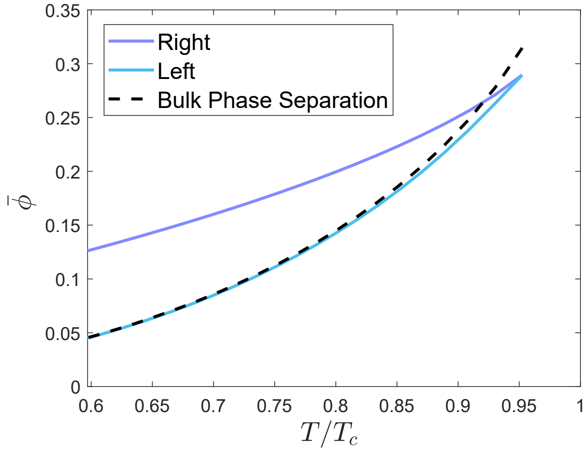


FIG. S4. $\bar{\phi}_{\text{right}}$ and $\bar{\phi}_{\text{left}}$ vs. $T/T_c$ when $L \to \infty$. The black-dashed line shows the saturation concentration for bulk phase separation $\phi_c$. $\bar{\phi}_{\text{left}}$ approaches $\phi_c$ in the low temperature limit.

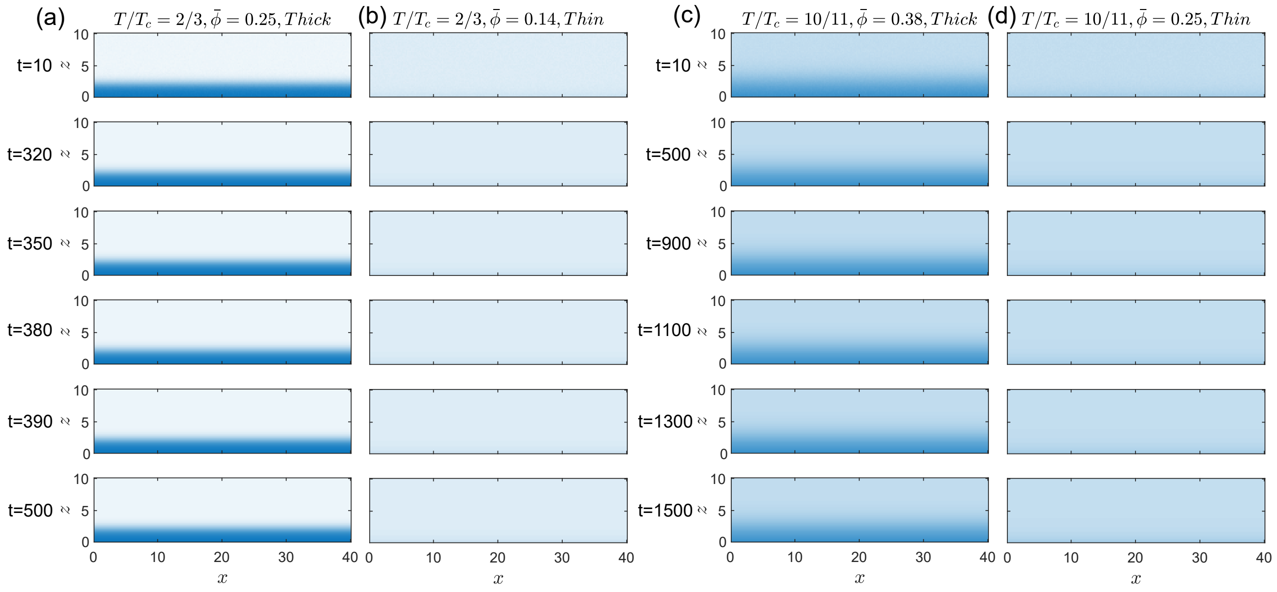


FIG. S5. Simulations of the two-dimensional systems below (a,b) and above (c,d) the wetting temperature. A periodic boundary condition is used in the $x$ direction with $d = 40$. At $t = 0$, a small perturbation is added to a homogeneous thick (a,c) or thin (b,d) profile. The interfaces in all four cases are stable.

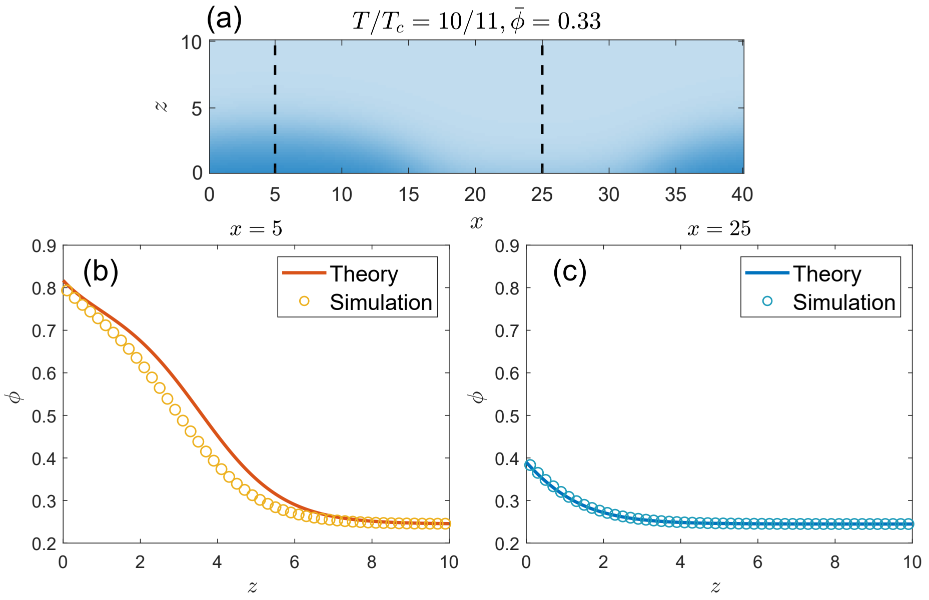


FIG. S6. (a) The simulated volume fraction distribution for $T > T_w$, corresponding to the prewetting state. The chemical potential in this system is $\mu = -0.0037$, with which we can calculate the volume-fraction profiles of the thick and thin states numerically. (b,c) The volume-fraction profiles of the droplet region (b) and the non-droplet region (c) in the $z$ direction are close to the predicted ones. The black dashed lines in (a) indicate where we take the simulated profiles.

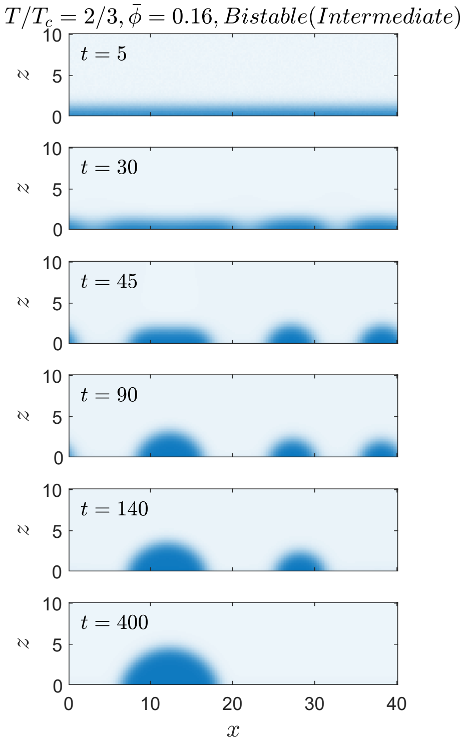


FIG. S7. Simulation of a two-dimensional system below $T_{sc}$. This system is in the lower part of the bistable region, where an intermediate solution and a thin solution coexist. At $t = 0$, a small perturbation is added to a homogeneous intermediate profile. The interface is unstable and evolves into a partial-wetting state.
